# Tomato fruit susceptibility to fungal disease can be uncoupled from ripening by suppressing susceptibility factors

**DOI:** 10.1101/2020.06.03.132829

**Authors:** Christian J. Silva, Casper van den Abeele, Isabel Ortega-Salazar, Victor Papin, Jaclyn A. Adaskaveg, Duoduo Wang, Clare L. Casteel, Graham B. Seymour, Barbara Blanco-Ulate

## Abstract

The increased susceptibility of ripe fruit to fungal pathogens poses a substantial threat to crop production and marketability. Here, we coupled transcriptomic analyses with mutant studies to uncover critical processes associated with defenses and susceptibility in tomato (*Solanum lycopersicum*) fruit. Using unripe and ripe fruit inoculated with three fungal pathogens, we identified common pathogen responses reliant on chitinases, WRKY transcription factors, and reactive oxygen species detoxification. We established that the magnitude and diversity of defense responses do not significantly impact the interaction outcome, as susceptible ripe fruit mounted a strong defense response to pathogen infection. Then, to distinguish features of ripening that may be responsible for susceptibility, we utilized non-ripening tomato mutants that displayed different susceptibility patterns to fungal infection. Based on transcriptional and hormone profiling, susceptible tomato genotypes had losses in the maintenance of cellular redox homeostasis, while jasmonic acid accumulation and signaling coincided with defense activation in resistant fruit. We identified and validated a susceptibility factor, pectate lyase (*PL*). CRISPR-based knockouts of *PL*, but not polygalacturonase (*PG2a*), reduced susceptibility of ripe fruit by >50%. This study suggests that targeting specific genes that drive susceptibility is a viable strategy to improve the resistance of tomato fruit against fungal disease.

**Highlight:** Increased susceptibility to fungal disease during tomato ripening is driven by the accumulation of susceptibility factors and not the lack of defense responses.

## Introduction

Half of all fruits and vegetables produced globally are lost each year (Gustavsson *et al*., 2011). While the causes of losses vary by region and commodity, fungal phytopathogens have a widespread role, as 20-25% of all harvested fruits and vegetables are lost to rotting caused by such fungi (Sharma *et al*., 2009). In fleshy fruit, this issue is exacerbated because, in general, fruit become more susceptible to fungal pathogens as they ripen (Prusky, 1996; Blanco-Ulate *et al*., 2016). Ripening-associated susceptibility has been demonstrated in multiple commodities including climacteric fruit such as tomato, stone fruit, banana, apple, and pear, as well as non-climacteric fruit such as strawberry, cantaloupe, citrus, and pineapple (Zhang *et al*., 1999; Gell *et al*., 2008; Morales *et al*., 2008; Cantu *et al*., 2009; Lassois *et al*., 2010; Chiu *et al*., 2013; Alkan *et al*., 2015; Petrasch *et al*., 2019; Lafuente *et al*., 2019; Barral *et al*., 2019).

The most devastating postharvest pathogens in fruit are those with necrotrophic lifestyles, which deliberately kill host tissue, resulting in rotting. Example pathogens include the model necrotrophic fungi *Botrytis cinerea* and *Sclerotinia sclerotiorum* as well as *Monilinia* spp., *Alternaria* spp., *Rhizopus* spp., *Penicillium* spp., and *Fusarium* spp. (Nunes, 2012; Bautista-Baños, 2014; van Kan *et al*., 2014; Liang and Rollins, 2018; Petrasch *et al*., 2019). Plant defense responses against necrotrophic fungi are multi-layered, involving (1) recognition of pathogen-associated molecular patterns (PAMPs), such as chitin or chitosan, by pattern recognition receptors (PRRs), (2) intracellular signaling through mitogen-activated protein (MAP) kinase cascades, (3) induction of downstream defenses by coordinated activity of phytohormones, particularly ethylene and jasmonic acid (JA), (4) cell wall fortifications, and (5) production of various secondary metabolites and antifungal proteins (van der Ent and Pieterse, 2012; Mbengue *et al*., 2016; Pandey *et al*., 2016; AbuQamar *et al*., 2017; Veloso and van Kan, 2018). However, most defense strategies have been studied in leaves, and their utilization and effectiveness in fruit have been assessed only with single pathogens (Cantu *et al*., 2009; Alkan *et al*., 2015; Ahmadi-Afzadi *et al*., 2018).

The outcome of any fruit-necrotroph interaction relies on the balance between the presence or induction of defenses and the contributions of susceptibility factors. Though induced defenses are heavily studied in plant immunity, the impact of preformed (or ‘constitutive’) defenses and susceptibility factors are less researched (van Schie and Takken, 2014). Preformed defenses include structural barriers, such as the cell wall and cuticle, and the accumulation of secondary metabolites (Wittstock and Gershenzon, 2002; Veronese *et al*., 2003), while susceptibility factors include the abundance of simple sugars and organic acids or activity of host cell wall modifying proteins (Cantu *et al*., 2008; Centeno *et al*., 2011). A sufficient understanding of ripening-associated susceptibility requires a characterization of the ripening program’s impact on (1) the ability of the host to express necessary defense genes upon pathogen challenge, (2) the integrity of preformed defenses, and (3) the abundance of susceptibility factors.

In this study, we first applied a transcriptomic approach to characterize core tomato fruit responses to three fungal pathogens and changes in gene expression that occur during ripening to promote susceptibility. To identify core responses that are not merely pathogen-specific, we used three pathogens with necrotrophic infection strategies: *B. cinerea, Rhizopus stolonifer*, and *Fusarium acuminatum*. Using well-established defense gene classifications, we developed profiles of host defense gene expression responses in unripe and ripe fruit. We then determined the susceptibility phenotypes of three non-ripening mutants: *Colorless non-ripening* (*Cnr*), *ripening inhibitor* (*rin*), and *non-ripening* (*nor*), which have unique defects in ripening features (Vrebalov *et al*., 2002; Giovannoni *et al*., 2004; Manning *et al*., 2006; Ito *et al*., 2017; Wang *et al*., 2019*b*; Gao *et al*., 2019, 2020). After demonstrating that each mutant has distinct susceptibility to disease, we identified ripening genes whose expression changes may impact the disease outcome. By integrating our transcriptomic data and mutant analyses, we found preformed defenses and susceptibility factor candidates associated with *B. cinerea* infections. Using CRISPR-based mutants, we established that one candidate, the pectin-degrading enzyme pectate lyase is indeed a disease susceptibility factor in tomato fruit.

## Materials and methods

### Plant material

Tomato (*Solanum lycopersicum*) *c*.*v*. ‘Ailsa Craig’ (AC), isogenic non-ripening mutants *rin, nor*, and *Cnr*, and CRISPR-based *PL* (PL5-4) and *PG2a* (PG21) mutants with azygous control plants (Wang *et al*., 2019*a*) were grown under standard field conditions in the Department of Plant Sciences Field Facilities at the University of California, Davis. Fruit were tagged at three days post-anthesis (dpa) and harvested at 31 dpa for mature green (MG) and at 42 dpa for red ripe (RR) or equivalent for ripening mutants. Ripening stages were confirmed by the color, size, and texture of the fruit.

The CRISPR line genotypes were confirmed by PCR of DNA prepared from leaf punches using Thermo Scientific Phire Plant Direct PCR Kits (Thermo Fisher Scientific, Waltham, MA, USA). Genes were amplified using the Phire PCR enzyme in thermocycler conditions: 98 °C denaturation for 5 min; 35 cycles of 98 °C for 5 s, 56 °C for 5 s, 72 °C for 20 s; and a final 1 min extension of 72 °C. The PCR products were column purified and Sanger sequenced. The primers were CAAAGGAATAGTATTCTCCTTCTC/CAGTTCCATGGAAAATGACTTTC for PG21 and GTGGTACCGGAAATCCAATC/CAATGATCCACCCAAACATG for PL5-4.

### Fungal culture and fruit inoculation

*R. stolonifer* and *F. acuminatum* isolates were taken from infections of produce and identified through morphological and sequencing methods (Petrasch *et al*., 2019). *B. cinerea* (B05.10), *R. stolonifer*, and *F. acuminatum* cultures were grown on 1% potato dextrose agar media. Conidia were harvested from sporulating cultures in 0.01% TWEEN⍰ 20 (Sigma-Aldrich Corporation, USA) and counted. Spore suspensions were stored for less than a month at −20 °C until use. Immediately prior to inoculation, spores were diluted with sterile milli-Q water to 500 conidia/μL, 30 conidia/μL, or 1000 conidia/μL for *B. cinerea, R. stolonifer*, and *F. acuminatum*, respectively. Fruit were surface disinfected by dipping twice in a 10% NaOCl solution for 30 s and followed by a deionized water wash. The blossom end halves of fungal-inoculated and wounded fruit were punctured to ca. 2 mm depth and ca. 1 mm diameter using a sterile micropipette tip. Each fruit used to measure disease incidence and severity was punctured at six sites; each fruit used for RNA extraction and transcriptomic analysis was punctured at 15 sites. For inoculated fruit, each puncture site was inoculated with 10 μL of spore solution, while no inoculum was introduced at puncture sites on wounded fruit. Healthy controls were not wounded or inoculated. Fruit were incubated for up to three days at 25 °C in high-humidity containers. Each biological replicate of each treatment (i.e. combination of genotype, ripening stage, and infection status) consisted of approximately eight fruit. Five biological replicates of each treatment were used for transcriptomic analysis, and four biological replicates of each treatment were used for measurements of disease progression.

### Disease incidence and severity measurements

Fruit disease incidence and severity were measured at 1, 2, and 3 days post-inoculation (dpi). Disease incidence was the percentage of inoculated sites displaying visual signs of tissue maceration or soft rot. Disease severity was calculated as the average lesion diameter (in mm) of each inoculation site displaying signs of rot.

### RNA extraction and library preparation

At 1 dpi, fruit pericarp and epidermal tissue of the blossom end halves of healthy, wounded, and infected fruit were collected and immediately frozen in liquid nitrogen and lysed using a Retsch® Mixer Mill MM 400 (Retsch, Germany). RNA was extracted from 1 gram of ground material as described in Blanco-Ulate *et al*., 2013. The purity and concentration of the extracted RNA were determined with a NanoDrop One Spectrophotometer (Thermo Scientific, USA) and a precise concentration measurement with the Qubit 3 (Invitrogen, USA). The integrity of the RNA was confirmed via agarose gel electrophoresis.

126 cDNA libraries were prepared using the Illumina TruSeq RNA Sample Preparation Kit v.2 (Illumina, USA) from isolated RNA. Each library was barcoded and analyzed with the High Sensitivity DNA Analysis Kit for the Agilent 2100 Bioanalyzer (Agilent Technologies, USA). Libraries were sequenced as single-end 50-bp reads on an Illumina HiSeq 4000 platform by the DNA Technologies Core at the UC Davis Genome Center.

### RNA sequencing and data processing

Raw sequencing reads were trimmed for quality and adapter sequences using Trimmomatic v0.33 (Bolger *et al*., 2014) with the following parameters: maximum seed mismatches = 2, palindrome clip threshold = 30, simple clip threshold = 10, minimum leading quality = 3, minimum trailing quality = 3, window size = 4, required quality = 15, and minimum length = 36. Trimmed reads were mapped using Bowtie2 (Langmead and Salzberg, 2012) to combined transcriptomes of tomato (SL4.0 release; http://solgenomics.net) and one of the three pathogens: *B. cinerea* (http://fungi.ensembl.org/Botrytis_cinerea/Info/Index), *F. acuminatum* (Petrasch *et al*., 2019), or *R. stolonifer* (Petrasch *et al*., 2019). Count matrices were made from the Bowtie2 results using sam2counts.py v0.91 (https://github.com/vsbuffalo/sam2counts/). Only reads that mapped to the tomato transcriptome were used in the following analyses. A summary of the read mapping results can be found in **Supplemental Table S1**. The datasets for this study have been deposited in the Gene Expression Omnibus (GEO) database under the accession GSE148217.

### Differential expression analysis

The Bioconductor package DESeq2 (Love *et al*., 2014) was used to perform normalization of read counts and differential expression analyses for various treatment comparisons. Differentially expressed (DE) genes for each comparison were those with an adjusted p-value of less than or equal to 0.05.

### Functional annotation and enrichment analyses

Gene Ontology (GO) terms were retrieved from SolGenomics. Annotations for transcription factors and kinases were generated using the automatic annotation tool from iTAK (Zheng *et al*., 2016). NBS-LRR family members were identified from Andolfo et al., 2014. Kyoto Encyclopedia of Genes and Genomes (KEGG) annotations were determined using the KEGG Automatic Annotation Server (Moriya *et al*., 2007), and hormone annotations were derived from these (**Supplemental Table S1**). GO enrichments were performed with the goseq package in R (Young *et al*., 2010), while enrichments for all other annotations were performed using a Fisher test with resulting p-values adjusted via the Benjamini and Hochberg method (Benjamini and Hochberg, 1995).

### Measurement of phytohormones

Ethylene emission was measured in MG and RR fruit from the day of harvest through 3 dpi. Headspace gas (3 ml) from weighed fruit in sealed 1-L containers was extracted after 30 minutes in a Shimadzu CG-8A gas chromatograph (Shimadzu Scientific Instruments, Kyoto, Japan). Sample peaks were measured against an ethylene standard of 1 ppm. Ethylene production was calculated from the peak height, fruit mass, and incubation time.

JA was measured using liquid chromatography coupled to tandem mass spectrometry and internal standards as in Patton *et al*., 2020 with modifications. Briefly, frozen tissue was lyophilized, weighed and extracted in isopropanol:H2O:HCL_1MOL_(2:1:0.005) with 100 μl of internal standard solution (1000 pg) as previously described (Casteel *et al*., 2015). Samples were evaporated to dryness, resuspended in 100 μl of MeOH, filtered, and 10 µl samples injected into an Agilent Technologies 6420 Triple Quad Liquid Chromatography-Tandem Mass Spectrometry instrument (Agilent, USA). A Zorbax Extend-C18 column 3.0×150mm (Agilent, USA) with 0.1% formic acid in water (A) and 0.1% (v/v) formic acid in acetonitrile (B) at a flow rate of 600 mL min^−1^ was used. The gradient was 0–1 min, 20% B; 1–10 min, linear gradient to 100% B; 10-13 min, 100% A.

## Results

### Susceptibility of tomato fruit to fungal infections by *Botrytis cinerea, Fusarium acuminatum*, and *Rhizopus stolonifer* increases during ripening

To characterize tomato fruit responses to fungal infection at unripe (MG) and ripe (RR) stages, we inoculated fruit (*c*.*v*. ‘Ailsa Craig’) with *B. cinerea, F. acuminatum*, or *R. stolonifer* spores. Each pathogen successfully infected RR fruit, producing visible water-soaked lesions and mycelial growth by 3 dpi, whereas MG fruit remained resistant and, except in samples inoculated with *R. stolonifer*, had a dark, necrotic ring around the inoculation sites (**Fig. 1A**), a feature of pathogen response that did not appear in wounded fruit. Thus, MG fruit resistance and RR fruit susceptibility are a feature common to multiple necrotrophic infections. We hypothesized that these susceptibility phenotypes are the result of (1) differences in defense responses at each ripening stage and (2) developmental processes during ripening that alter the levels of preformed defenses and susceptibility factors (**Fig. 1B**). First, we assumed that, compared to a robust defense response in MG fruit, RR fruit have a weaker response, consisting of fewer genes induced, less diverse functionality, and absent expression of critical genes. Additionally, we predicted that ripening may decrease the expression of preformed defenses and increase the expression of susceptibility factors, which create a more favorable environment for infection.

**Fig. 1:**
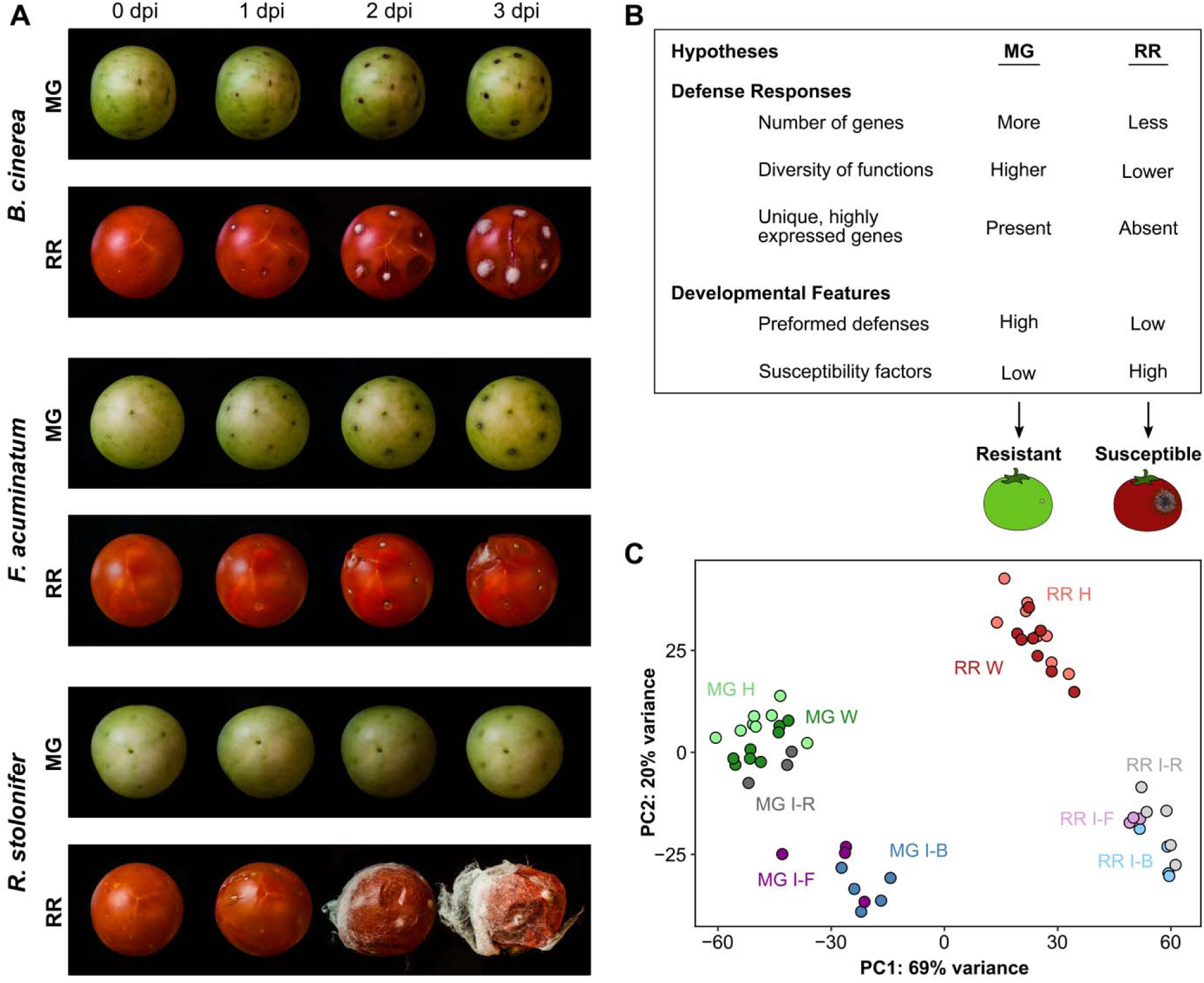
Tomato fruit responses to *B. cinerea, F. acuminatum*, and *R. stolonifer*. **(A)** Disease progression in inoculated mature green (MG) and red ripe (RR) fruit each day up to 3 days post-inoculation (dpi). **(B)** Hypotheses of why MG fruit are resistant while RR fruit are susceptible to fungal disease. **(C)** Principal component analysis of total mapped RNA-Seq tomato reads. Color corresponds to treatment. H = healthy, W = wounded, I = inoculated 1 dpi, B = *B. cinerea*, F = *F. acuminatum*, R = *R. stolonifer*.

### Susceptible ripe fruit respond to pathogens with a larger, more diverse set of defense genes than resistant unripe fruit

To test if defense responses to fungal pathogens are compromised in RR compared to MG fruit, we sequenced mRNA from *B. cinerea-, F. acuminatum-*, and *R. stolonifer-*inoculated fruit at 1 dpi, an early timepoint at which either a resistant or susceptible phenotype becomes apparent. We included healthy and wounded MG and RR fruit from the same timepoint as controls. A principal component analysis (PCA) of the mapped normalized reads for all tomato genes (**Fig. 1C**) revealed that the major driver separating sample data was the ripening stage (PC1, 69%), while inoculation status accounted for less of the separation (PC2, 20%). The one exception to this pattern was the *R. stolonifer*-inoculated MG samples, which clustered with the healthy and wounded MG samples, suggesting that unripe fruit did not display strong responses to this pathogen and yet remained resistant. However, quantification of normalized pathogen reads (**Supplemental Fig. S1A**) confirmed that all three pathogens were detectable at 1 dpi even in MG samples.

To identify the responses for each ripening stage common to all three pathogens, we performed a differential expression analysis between inoculated and healthy samples for MG and RR fruit. We chose the healthy samples as controls for these comparisons in order to capture responses to necrotrophic infection which may share features with mechanical wounding. Of all 34,075 protein-coding genes found in the tomato transcriptome, 9,366 (27.5%) were found to be differentially expressed (*P*_*ad*j_ < 0.05) in response to inoculation in fruit at 1 dpi in at least one comparison (**Supplemental Table S2**). Of these, 475 genes were significantly upregulated in MG fruit in response to all three pathogens, corresponding to the MG core response (**Fig. 2A**), whereas 1,538 genes formed the RR core response (**Fig. 2B**). The MG core response overlapped substantially with the wounding response in MG fruit (**Supplemental Fig. S1B**), which suggests that unripe fruit activate similar functions when responding to pathogen attack and mechanical damage. However, this large overlap is also due to the similarity between the gene expression profiles of wounded and *R. stolonifer*-inoculated samples as seen in the PCA (**Fig. 1C**). In contrast, the lack of a strong wounding response in RR fruit indicates that nearly all RR core response genes were strictly pathogen-related (**Supplemental Fig. 1B**). Downregulated genes in response to infection were largely unique to each pathogen, with only 57 and 225 downregulated across all three pathogens in MG and RR fruit, respectively, and thus we decided to continue our analysis only on the upregulated core response genes. Complete lists of gene set intersections of upregulated and downregulated genes are in **Supplemental Table S3**.

**Fig. 2:**
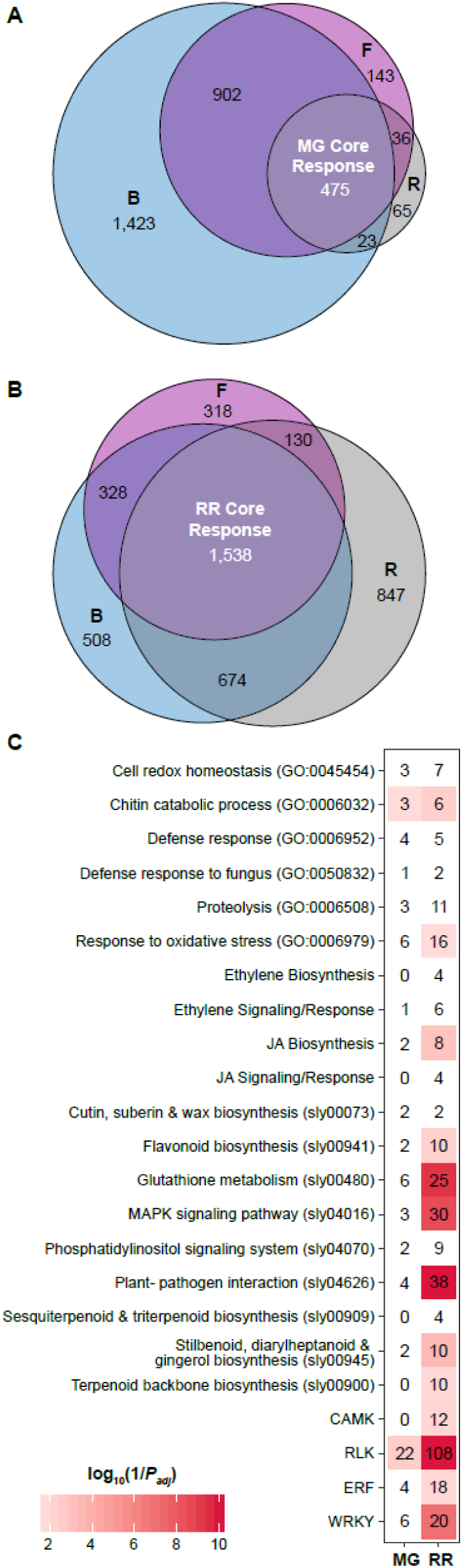
Tomato core responses to fungal inoculations. **(A-B)** Euler diagram of tomato genes upregulated in response to inoculation in (**A**) mature green (MG) or (**B**) red ripe (RR) fruit. B = *B. cinerea*, F = *F. acuminatum*, R = *R. stolonifer*. Core responses are shown in white. (**C**) Enrichments of various defense-related classes in the MG and RR core responses. The scale is the log_10_(1/P_adj_). Values greater than 10 were converted to 10 for scaling purposes. Numbers in each tile indicate the number of genes within each classification. JA = jasmonic acid, MAPK = mitogen-activated protein kinase, CAMK = calmodulin-dependent protein kinase, RLK = receptor-like kinase, ERF = ethylene responsive factor.

We then assessed the MG and RR core responses for the presence of various well-established gene classifications related to pathogen defense, including selected GO terms, KEGG pathways, transcription factor (TF) families, hormone biosynthesis, signaling, and response genes, and receptor-like kinase (RLK) genes (**Fig. 2C**). For each category, we performed enrichment analyses (*P*_adj_ < 0.05) to identify classifications of particular importance in both MG and RR core responses. A total of 70 defense genes were identified in the MG core response. Interestingly, these were enriched in only two categories: chitin catabolic process (GO:0006032) and RLK genes. The RR core response was enriched in 13 defense categories, including the plant-pathogen interaction (sly04626) and MAPK signaling pathways (sly4016), secondary metabolite biosynthesis pathways (sly00900, sly00941, sly00945), WRKY and ERF (ethylene responsive factor) transcription factors, RLKs, and JA biosynthesis. Altogether, 302 defense genes were identified among the RR core response. Thus, in contrast to their respective susceptibility phenotypes, RR fruit appear to mount a more robust and diverse defense response than MG fruit early during inoculation, demonstrating that, contrary to our initial hypothesis, weakened defense responses in RR fruit are not a contributor to ripening-associated susceptibility.

However, it is possible that tomato fruit resistance to necrotrophs could be determined by a small number of genes that were exclusive to the MG core response. Out of the 70 defense genes in the MG core response, 27 were not found in the RR core response (**Fig. 3**). These 27 genes are heterogeneous, representing 12 different defense categories. Notable genes in this category include a three-gene cluster of *PR-10* family proteins (GO:0006952), a chitinase previously identified during infections of tomato with *Cladiosporum fulvum* (*Solyc10g055810*, Danhash et al., 1993), and an ERF active at the onset of ripening (Liu *et al*., 2015*b*). Although these 27 genes were not in the RR core response, most of them were induced during RR infections by one or two of the pathogens studied. Only seven were not upregulated by any of the three pathogens in RR fruit, including the ERF mentioned above (*Solyc03g118190*), as well as three RLK genes, two glutaredoxin genes involved in the response to oxidative stress, and a cysteine protease. Given that each of these genes belongs to a large family of genes whose members are often functionally redundant, and their average expression levels in infected MG fruit were fairly low (normalized read counts 8.13 – 149.07), we consider it unlikely that the lack of these genes in the RR core response contributes heavily to susceptibility.

**Fig. 3:**
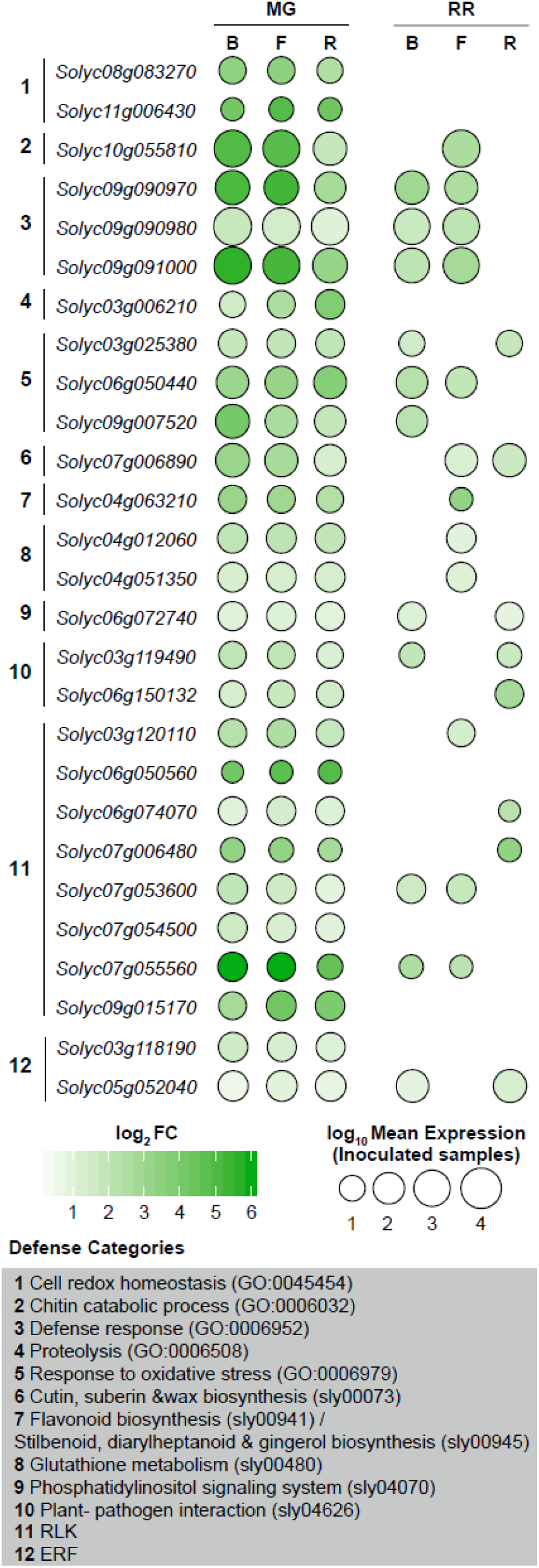
Defense genes in the mature green (MG) core response that are not in the red ripe (RR) core response. Dot sizes are proportional to the average normalized read count values from the inoculated fruit samples. RR= Red Ripe, B = *B. cinerea*, F = *F. acuminatum*, R = *R. stolonifer*, RLK = receptor-like kinase, ERF = ethylene responsive factor.

Additionally, the induction of defense genes in the RR core response could be ineffective if their expression levels were too weak compared to those seen in resistant MG fruit. We evaluated the levels of gene expression in inoculated RR fruit via a differential expression comparison (*P*_adj_ < 0.05) to inoculated MG fruit. Of all the RR core defense genes identified above, 269/302 (89.1%) are expressed at equal or greater levels (average log_2_FC = 2.16) in inoculated RR fruit compared to inoculated MG fruit for all three pathogens. Conversely, 33/302 (11.9%) of these defense genes were expressed at higher levels in MG fruit compared to RR fruit for at least one of the three pathogens (**Supplemental Table S4**). These genes are diverse, representing 15 different defense categories. Prominent genes in this category include *TAP1* (*Solyc02g079500*) and *TAP2* (*Solyc02g079510*), two peroxidases associated with defensive suberization in tomato (Roberts and Kolattukudy, 1989; Kesanakurti *et al*., 2012); *CHI3* (*Solyc02g082920*) and *CHI17* (*Solyc02g082930*) two chitinases associated with *C. fulvum* infection (Danhash *et al*., 1993), and the JA biosynthesis gene *OPR3* (*Solyc07g007870*). While it is possible that resistance may be determined by these genes, these results indicated that the differences in defense responses observed between MG and RR fruit are not likely solely responsible for differences in susceptibility, and, therefore, we considered the alternate hypothesis.

### Defects in regulation of ripening result in altered susceptibility to fungal infection

We explored the possibility that the increase in susceptibility to fungal pathogens is heavily influenced by a decline of preformed defenses and accumulation of susceptibility factors that occur during fruit ripening prior to pathogen challenge. To identify developmental features that are integral to fruit resistance or susceptibility, we utilized the isogenic non-ripening tomato mutants *Cnr, rin*, and *nor*, which produce fruit that lack most of the characteristic changes associated with normal ripening, such as color, texture, acidity, sugar accumulation, and ethylene production, but yet are phenotypically different from one another. All three mutant lines likely result from spontaneous gain-of-function mutations in transcription factors with key roles in the regulation of ripening (Vrebalov *et al*., 2002; Giovannoni *et al*., 2004; Manning *et al*., 2006; Ito *et al*., 2017; Wang *et al*., 2019*b*; Gao *et al*., 2019, 2020).

We inoculated fruit of these mutant genotypes at comparable stages to MG and RR wild-type fruit (i.e. “MG-like” and “RR-like”) with *B. cinerea, F. acuminatum*, and *R. stolonifer* and measured disease incidence and severity up to 3 dpi (**Fig. 4**). For all three pathogens at both MG-like and RR-like stages, only *nor* fruit were consistently resistant to infection. MG-like fruit of *Cnr* were the only unripe fruit susceptible to any pathogen, with both *B. cinerea* and *F. acuminatum* able to produce lesions on a significant number of these fruit. Consistent with this, *Cnr* RR-like were more susceptible than wild-type RR fruit to *B. cinerea*, with average disease severity (i.e. lesion size) nearly twice as great at 3 dpi (**Fig. 4A**). The fruit of *rin* at both MG-like and RR-like stages showed similar or slightly lower susceptibility to all pathogens when compared to wild-type, with the exception of a significant reduction in disease incidence to *F. acuminatum* at the RR-like stage. Because some ripening processes may promote susceptibility, others may maintain resistance, and others may have no impact, we hypothesized that the *Cnr, rin*, and *nor* mutations differentially affect ripening-associated genes or pathways that are critical to tip the balance towards either susceptibility or resistance.

**Fig. 4.**
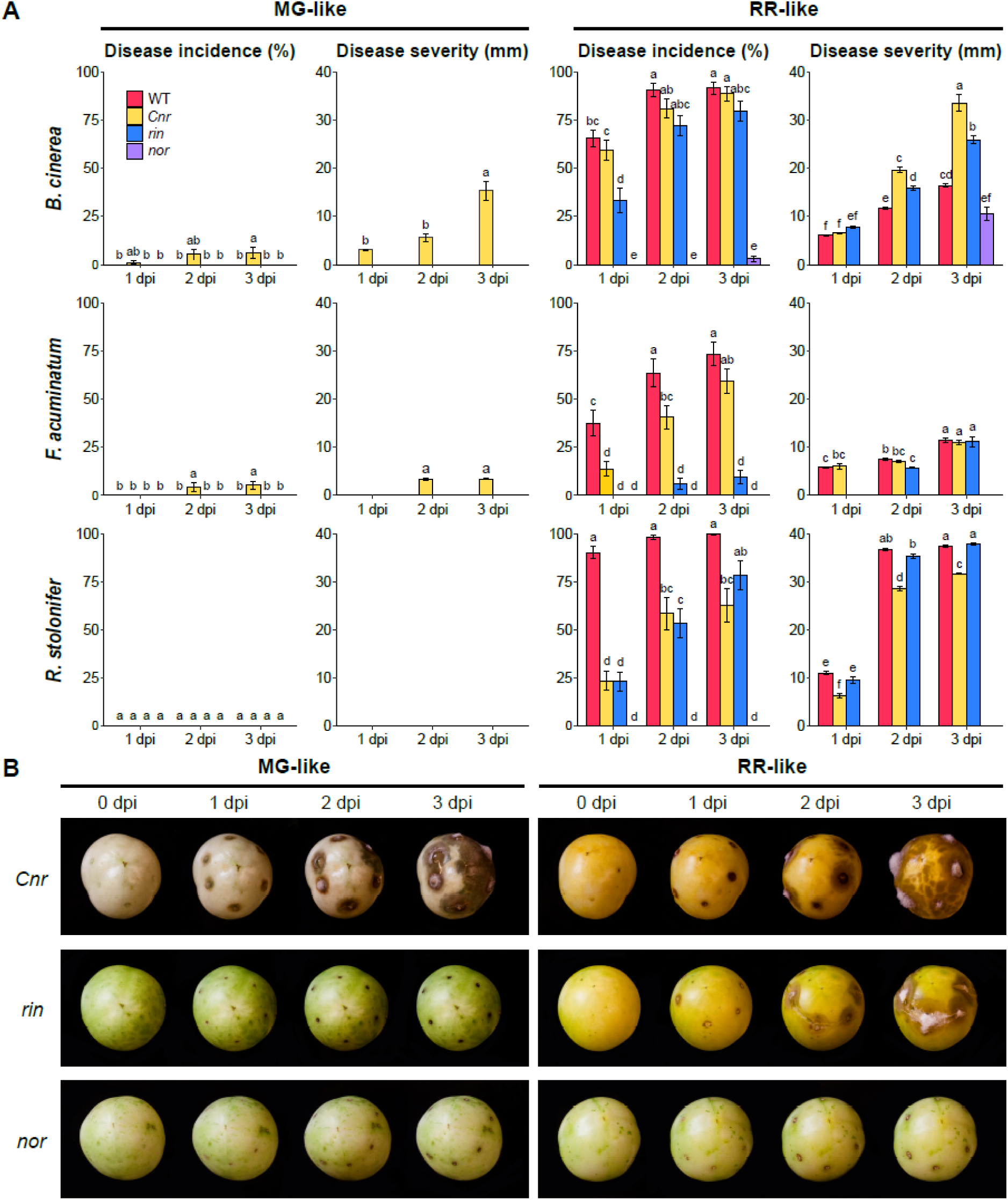
Susceptibility of the non-ripening mutants *Cnr, rin*, and *nor* to fungal infections. **(A)** Disease incidence and severity measurements for MG-like (left) and RR-like (right) fruit. Wild-type values are included for comparison. **(B)** Disease progression of *B. cinerea*-inoculated MG-like and RR-like fruit each day up to 3 days post-inoculation.

We sequenced mRNA from *B. cinerea*-inoculated and healthy fruit from the non-ripening mutants at MG-like and RR-like stages at 1 dpi. We chose *B. cinerea* inoculations because this pathogen showed the clearest differences in susceptibility phenotypes between these genotypes. We first characterized transcriptional responses of mutant fruit to pathogen challenge by using enrichment analysis of defense-related processes to determine if differences in defense responses could explain the distinct susceptibility phenotypes (**Supplemental Fig. 2A**). In most cases, the mutant fruit exhibited similar patterns of defense classification enrichments as wild-type fruit in both stages, with some notable exceptions. Compared to the other genotype-stage combinations, *Cnr* MG-like responses were deficient (i.e., less enriched) in the expression of genes from several prominent defense classifications, including chitin catabolic process (GO:0006032), the plant-pathogen interaction (sly04626) and glutathione metabolism (sly00480) pathways, ERF and WRKY transcription factors, and RLK and CAMK genes. Given that *Cnr* fruit were the only genotype at the MG-like stage to display susceptibility to *B. cinerea* infection, it can be suggested that these defense processes may be necessary for resistance in unripe fruit. However, these processes were enriched in the susceptible RR-like fruit of *Cnr* and *rin*, as well as wild-type RR fruit, which clearly indicates that they are not sufficient to result in a resistant outcome.

The role of ethylene and JA showed some variation amongst the mutants. For example, the responses of resistant *nor* fruit in both MG-like and RR-like fruit were noticeably less enriched in ethylene-associated pathways and more enriched in JA-associated pathways. These results suggest that JA-mediated defenses may contribute to tomato fruit resistance in the absence of ethylene, and that the *nor* mutation may activate JA-associated resistance. In support of this observation, levels of JA in healthy fruit appeared to be linked to resistance: they were highest in RR-like *nor* fruit, and only *nor* fruit experienced an increase in JA in the transition from MG-like to RR-like (**Supplemental Fig. 2B**). However, ethylene levels increase dramatically during ripening in wild-type fruit, but they remain low and even decrease slightly in both the susceptible *Cnr* and *rin* fruit as well as the resistant *nor* fruit (**Supplemental Fig. 2C)**. Still, ethylene biosynthesis is induced in all genotypes except *nor* in response to *B. cinerea* inoculation, and ethylene signaling/response genes are highly enriched in *Cnr* MG-like fruit (**Supplemental Fig. 2A**). Overall, with the exception of *Cnr* MG-like fruit, resistance or susceptibility in the non-ripening mutants cannot be merely explained by the presence and/or magnitude of defense responses.

### Fruit infections are promoted by a decrease in preformed defenses and an increase in insusceptibility factors during ripening

To identify genes that are involved in resistance or susceptibility that change during tomato fruit ripening, we used a differential expression analysis (*P*_adj_ < 0.05) comparing healthy RR/RR-like to healthy MG/MG-like fruit for each wild-type and mutant lines. In wild-type fruit, 6,574 genes were significantly downregulated in RR fruit compared to MG, while 5,674 genes were significantly upregulated (**Supplemental Table S2**). We used the susceptibility phenotypes and the transcriptional profiles of the mutant fruit to filter these ripening-associated genes and identify critical preformed defense mechanisms or susceptibility factors. Of the four genotypes, all except *nor* experience an increase in susceptibility in the transition from MG/MG-like to RR/RR-like fruit. Thus, we selected ripening-associated genes that showed the same expression pattern in wild-type, *Cnr* and/or *rin*, but not *nor*. This filtering resulted in 2,893 downregulated and 2,003 upregulated genes, respectively.

We assumed that effective preformed defenses will decrease during ripening. Thus, the set of filtered downregulated genes, being those that are highly expressed in healthy MG fruit compared to healthy RR fruit, should contain key genes related to preformed defenses. The filtered downregulated genes contained 251 defense genes, while upregulated genes included only 171 defense genes, indicating a net loss of about 80 genes in the transition from MG/MG-like to RR/RR-like susceptible fruit. Furthermore, the 251 defense genes from the filtered downregulated set were overrepresented by functional categories involved in reactive oxygen species (ROS) response and detoxification, proteolysis, and the biosynthesis of secondary metabolites (**Table 1**). These downregulated ROS-related genes spanned several subfamilies including thioredoxins, glutaredoxins, glutathione S-transferases, and peroxidases. Among the downregulated proteolytic genes were several subtilisin-like proteases, including *SBT3* (*Solyc01g087850*; Meyer et al., 2016). Lastly, in addition to several genes involved the methylerythritol 4-phosphate (MEP) pathway of terpenoid biosynthesis, two copies of the lignin biosynthesis gene *CCoAOMT* (*Solyc01g107910, Solyc04g063210*) were also among the filtered downregulated class, suggesting that cell wall fortification could be inhibited upon infection. These results indicate that ripening involves a loss of multiple defense genes, and that the preexisting levels of genes involved in ROS regulation, proteolysis, and secondary metabolite biosynthesis may be critical for resistance.

**Table 1.** Defense categories enriched in a subset of significantly downregulated genes during ripening of healthy tomato fruit. The significance cut-off for the enrichments is *P*_adj_ < 0.05. Full enrichment results for both upregulated and downregulated defense genes can be found in **Supplemental Table S5**.

| Defense Category | Number of Genes | Example Functions |
| --- | --- | --- |
| Cell redox homeostasis (GO:0045454) | 24 | Thioredoxins, glutaredoxins |
| Defense response (GO:0006952) | 6 | MLO-like proteins, Sn-1 proteins |
| Proteolysis (GO:0006508) | 36 | Subtilisin-like proteases (SBT2, SBT3) |
| Response to oxidative stress (GO:0006979) | 16 | Peroxidases |
| Flavonoid biosynthesis (sly00941) | 5 | Caffeoyl-CoA O-methyltransferase |
| Glutathione metabolism (sly00480) | 18 | Glutathione S-transferases |
| MAPK signaling pathway (sly04016) | 17 | Protein phosphatase 2C, RBOH proteins |
| Phosphatidylinositol signaling system (sly04070) | 5 | Phosphatidylinositol phospholipase C |
| Plant-pathogen interaction (sly04626) | 15 | Disease resistance protein RPM1 |
| Terpenoid backbone biosynthesis (sly00900) | 8 | Geranylgeranyl diphosphate synthase |
| CAMK | 8 | Calcium-dependent kinases |
| RLK | 78 | Lectin receptor kinases, Leucine-rich repeat kinases |
| ERF | 8 | ERFA2, ERFC2, ERFC3 |

Finally, we evaluated filtered upregulated genes that are highly expressed in healthy RR fruit compared to healthy MG fruit, as they may include potential susceptibility factors. Since there is little scientific literature on classes of genes that may constitute susceptibility factors in plants, we focused on the upregulated genes that were highly expressed in the RR/RR-like fruit of the susceptible genotypes. Such genes may have disproportionate impacts on susceptibility due to their high expression. To identify these genes, we calculated average normalized read count values for each gene across WT, *Cnr*, and *rin* RR/RR-like fruit. The distribution of these values over the filtered upregulated genes is a notably long-tailed distribution with a range of 2.43 to 179,649.29 and an average of 1,295. We identified genes with abnormally high expression values by selecting outliers (i.e., values above 1.5 * the inter-quartile range) from a log_10_-transformed distribution of the data. This resulted in a list of 16 genes (**Table 2**). They include several genes previously discovered to be active during tomato fruit ripening, including the flavor volatile biosynthesis gene *ADH2* (*Solyc06g059740*; Speirs et al., 1998), the carotenoid biosynthesis gene *Z-ISO* (*Solyc12g098710*; Fantini et al., 2013), the pectin-degrading enzymes *PG2a* (*Solyc10g080210*; Sheehy et al., 1987) and *PL* (*Solyc03g111690*; Uluisik et al., 2016), and the ethylene receptor *ETR4* (*Solyc06g053710*; Tieman and Klee, 1999), among other genes involved in carbohydrate metabolism.

**Table 2.** Highly expressed genes in susceptible RR/RR-like fruit. Names and ripening functions were determined via BLAST and literature searches.

| **Accession** | **Average RR/RR-like Expression** | **Name** | **Ripening Function** |
| --- | --- | --- | --- |
| *Solyc06g059740* | 99,772.18 | *SIADH2* | Flavor aldehyde biosynthesis |
| *Solyc08g065610* | 64,989.08 | *SIVPE3* | Sugar metabolism |
| *Solyc03g111690* | 25,643.87 | *SIPL* | Pectin degradation |
| *Solyc10g080210* | 25,044.06 | *SIPG2a* | Pectin degradation |
| *Solyc08g014130* | 21,514.72 | *SIIPMS2* | Unknown |
| *Solyc10g076510* | 20,051.40 | -- | Unknown |
| *Solyc07g047800* | 19,462.21 | -- | Unknown |
| *Solyc12g005860* | 19,048.01 | -- | Unknown |
| *Solyc08g080640* | 17,227.90 | *SINP24* | Unknown |
| *Solyc12g098710* | 15,070.45 | *SIZ-ISO* | Carotenoid biosynthesis |
| *Solyc09g009260* | 14,572.63 | *SIFBA7* | Sugar metabolism |
| *Solyc10g024420* | 14,103.56 | -- | Unknown |

While any of these genes has the potential to impact susceptibility, cell wall-degrading enzymes such as *PL* and *PG2a*, which facilitate fruit softening during ripening, represent especially good candidates given both the importance of cell wall integrity in defense against fungal pathogens and previous research on RNAi-developed mutants in tomato (Cantu et al., 2008; Yang et al., 2017). To validate the impact of *PG2a* and *PL* expression in wild-type RR fruit on susceptibility to *B. cinerea*, we utilized CRISPR-based mutants in each of these genes (Wang *et al*., 2019*a*). RR fruit of the CRISPR-PL line, but not the CRISPR-PG2a mutant, demonstrated reduced susceptibility to *B. cinerea* compared to the azygous WT control line (**Fig. 5**). At 3 dpi, disease incidence in the PL lines was 56% lower than azygous lines. We concluded that the ripening-associated pectate lyase enzyme is a major susceptibility factor for *B. cinerea* infection in tomato fruit.

**Fig. 5.**
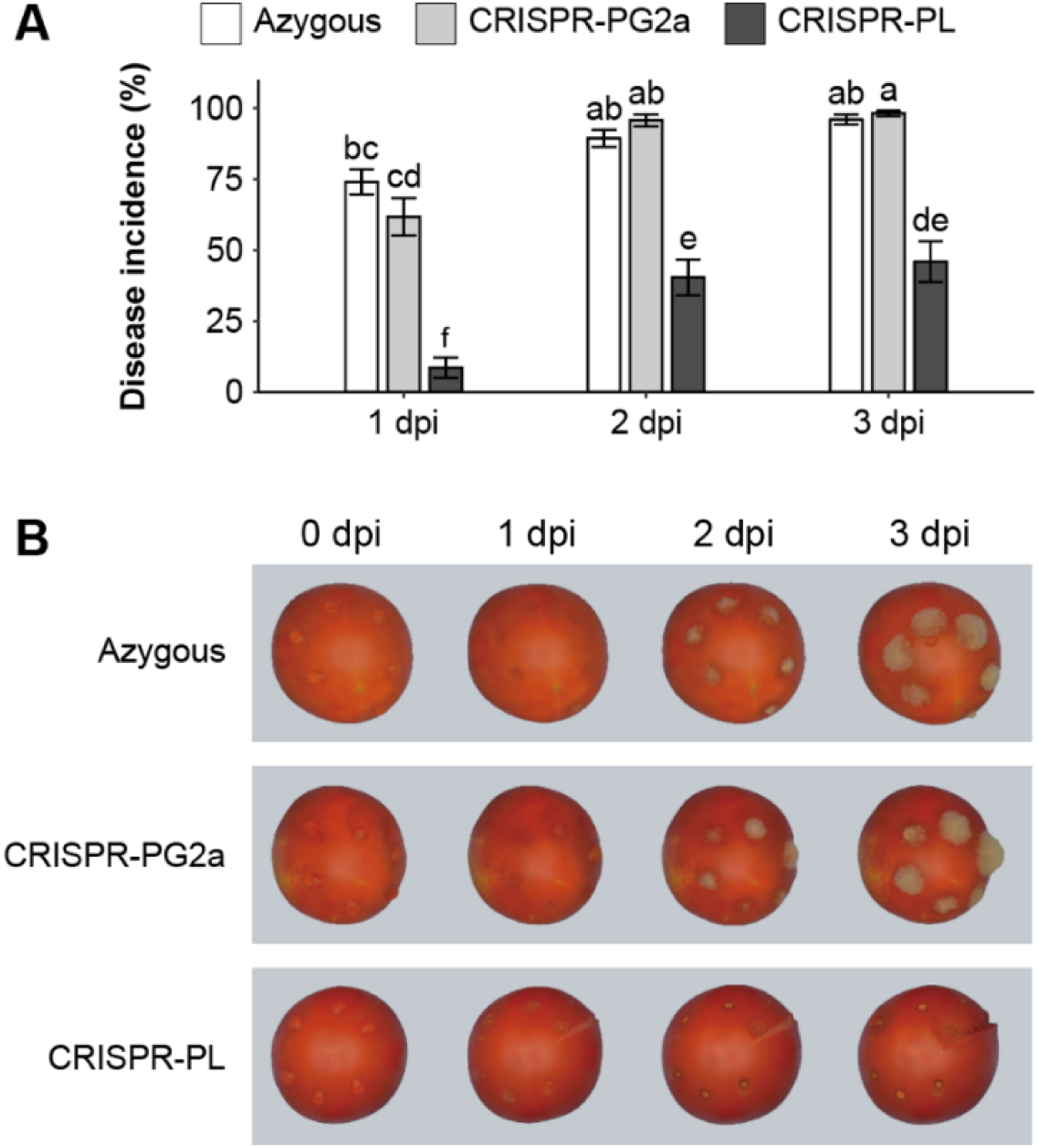
Inoculations of CRISPR lines with *Botrytis cinerea*. **(A)** Disease incidence measurements at 1, 2, and 3 dpi. **(B)** Photos of representative inoculated tomatoes from 0 to 3 dpi.

## Discussion

Increased susceptibility to fungal pathogens during ripening is a feature of many fleshy fruit. During ripening, fruit may gradually lose either the ability to activate or the effectiveness of components of the plant immune system, defensive hormone production and signaling, and downstream transcriptional responses. Alternatively, ripening processes such as cell wall breakdown, simple sugar accumulation, changes in pH and secondary metabolite composition, and, in climacteric fruit, increased levels of ethylene, may impact the fruit’s capability to resist fungal attack (Prusky *et al*., 2013; Alkan and Fortes, 2015). The widespread nature of this phenomenon in diverse fruit pathosystems suggests that ripening-associated susceptibility is likely to be mediated by combinations of the above factors.

In tomato, ripening-associated susceptibility has been demonstrated not only for the model necrotrophic pathogen *Botrytis cinerea*, but for other fungal pathogens including *Colletotrichum gloeosporioides* (Alkan *et al*., 2015), *Rhizopus stolonifer*, and *Fusarium acuminatum* (Petrasch *et al*., 2019). Here, for the first time, we identified specific host responses in both resistant unripe (MG) and susceptible ripe (RR) fruit that are common to multiple pathogens and thus represent core responses to fungal infection. Most prominently, these core responses featured RLKs, WRKY and ERF transcription factors, JA biosynthesis, and chitin catabolism. Some genes that appear in both the MG core and RR core responses were previously-studied components of plant immunity in tomato, including the JA biosynthesis gene *LoxD* (Yan *et al*., 2013), the subtilisin-like protease *SBT3* (Meyer *et al*., 2016), the peroxidase *CEVI-1* (Mayda *et al*., 2000), and the chitinase *CHI9* (Danhash *et al*., 1993).

However, most defense genes uncovered were found solely in the RR core response. These included several well-known defense genes that were only expressed in RR fruit, such as *WRKY33* (Liu *et al*., 2015*a*), the ERF *PTI5* (He *et al*., 2001; Gu *et al*., 2002; Wu *et al*., 2015), the RLK *TPK1b* (AbuQamar *et al*., 2008), and the MAP kinase *MPK3* (Kandoth *et al*., 2007; Stulemeijer *et al*., 2007; Zhang *et al*., 2018). While the MG core response did contain some defense genes that were not found in the RR core response, expression of most of these genes was found in the RR response to one or two pathogens. Many of these were functionally similar to other RR core response genes, and were not expressed at high levels in inoculated fruit. Thus, the ability to quickly express a large amount of defense genes at high levels does not appear to be compromised in RR fruit, and ripening-associated susceptibility is therefore not sufficiently explained by differences in the diversity or magnitude of defense responses.

If defense responses do not determine the outcome of the interaction in tomato fruit, developmental features associated with ripening of healthy fruit may instead govern susceptibility. The highly complex transcriptional reprogramming during ripening allows for a large number of potential contributors to the increase in susceptibility. Ripening processes in tomato have been studied using non-ripening mutants such as *Cnr, rin*, and *nor*. In addition to being phenotypically distinct, these mutants display differential susceptibility patterns when inoculated with fungal pathogens. Previously, susceptibility to *B. cinerea* in tomato fruit was shown to be dependent on *NOR* but not *RIN*, though the role of *CNR* remained uncharacterized (Cantu *et al*., 2009). Our results with *B. cinerea* as well as *F. acuminatum* and *R. stolonifer* corroborate the roles of *NOR* and *RIN* while also proposing a role for *CNR* in tomato fruit defense against fungal pathogens. In addition to exhibiting hypersusceptibility to *B. cinerea* in RR-like fruit, *Cnr* MG-like fruit were the only fruit of this stage to exhibit any susceptibility. Unlike *rin* and *nor* fruit, *Cnr* fruit have altered cell wall architecture even in MG-like stages (Eriksson *et al*., 2004; Ordaz-Ortiz *et al*., 2009), a feature which may be exploited during fungal infection. Moreover, compared to all other fruit, *Cnr* MG-like fruit were deficient in their defense responses against *B. cinerea*. However, apart from *Cnr* MG-like fruit, differences in defense responses appeared to have little impact on susceptibility, as enriched defense categories were similar across both resistant and susceptible mutant fruit.

We took advantage of the susceptibility differences in the ripening mutants to unravel ripening components that may represent either declining preformed defenses or increasing susceptibility factors. Differential expression analyses carefully filtered based on susceptibility phenotypes revealed that several defense-related genes undergo changes in gene expression during the transition from MG/MG-like to RR/RR-like fruit. Most interestingly, declining preformed defenses appear to be overrepresented by gene categories involved in the mediation of ROS levels. Host regulation of ROS levels during early fungal infection is critical for both defense signaling and detoxification of ROS generated by the pathogen (Lehmann *et al*., 2015; Waszczak *et al*., 2018), and tomato fruit susceptibility to *B. cinerea* has been shown to be impacted by both of these roles. Improved resistance to *B. cinerea* in the ABA-deficient *sitiens* mutant has been shown to be the result of controlled ROS production, which promotes cell wall fortification (Asselbergh *et al*., 2007; Curvers *et al*., 2010), and a similar improved *B. cinerea* resistance is seen in tomato varieties genetically engineered to produce especially high amounts of antioxidant anthocyanins in fruit (Zhang *et al*., 2015). During ripening, losing control of ROS levels may thus represent the reduction of an important preformed defense.

Some features of ripening have the potential to be either a preformed defense or a susceptibility factor depending on the context. The ethylene burst that accompanies ripening in climacteric fruit is an example. Although ethylene is known for its involvement in defense against necrotrophs (van der Ent and Pieterse, 2012), its induction of the ripening program catalyzes downstream events that can be favorable for pathogen infections. Previous research suggests that inhibition of ethylene receptors in MG fruit can either increase or decrease resistance to *B. cinerea* depending on the concentration of inhibitor used (Blanco-Ulate *et al*., 2013). Thus, ethylene-mediated resistance may be dependent on careful regulation of ethylene levels, and the autocatalytic ethylene biosynthesis that occurs in wild-type fruit ripening may be detrimental. We observed that, although ethylene levels in healthy fruit did not correlate well with susceptibility, ethylene-related transcriptional responses were particularly prominent in susceptible fruit, especially *Cnr* MG-like. In addition to ethylene, JA is known to mediate resistance to necrotrophs in plants (Wasternack and Hause, 2013; Pandey *et al*., 2016). The enrichment of JA biosynthesis genes is seen in the RR core response, as well as the response to *B. cinerea* in all mutant fruit at both stages. Basal levels of JA in healthy fruit are highest in *nor* RR-like fruit, where they are nearly twice as high as levels in wild-type RR fruit. Moreover, *nor* fruit are the only fruit at which JA signaling/response genes are enriched in response to *B. cinerea* infection at both stages. Although JA is linked to the promotion of fruit ripening (Peña-Cortés *et al*., 2004), its role is much less prominent than ethylene, which may allow it to play a defense role in fruit without having the unintended consequence of promoting ripening and, in turn, susceptibility. However, the interplay between ethylene and JA and their impact of ripening-associated susceptibility requires further study.

Other features of ripening can increase susceptibility to fungal disease such as the disassembly of plant cell walls leading to fruit softening. Cell wall polysaccharide remodeling, breakdown, and solubilization in ripening fruit occurs as the result of various cell wall-degrading enzymes, particularly those that act on pectin (Brummell, 2006). The cell wall represents an important physical barrier to pathogen attack in plants (Malinovsky *et al*., 2014), and cell wall integrity and fortification improves tomato fruit resistance to *B. cinerea* infection (Cantu et al., 2008; Curvers et al., 2010). The enzymes PL and PG2a feature prominently in tomato fruit ripening and softening (Uluisik *et al*., 2016; Yang *et al*., 2017; Wang *et al*., 2019*a*) and accumulate in RR/RR-like fruit of susceptible genotypes. However, these enzymes do not have equal impact on fruit softening, as CRISPR-based mutants in *PL*, but not *PG2a*, result in a reduced rate of softening in RR fruit (Wang *et al*., 2019*a*). This differential impact on firmness is mirrored in the effect on susceptibility to *B. cinerea*, as the firmer CRISPR-PL mutant was less susceptible than both the CRISPR-PG2a mutant and the azygous control. Though RR fruit of the CRISPR-PG2a mutant did not exhibit increased *B. cinerea* resistance, PG2a may still contribute to susceptibility, as RNAi-mediated knockdown of *PG2a* together with the expansin gene *Exp1* increases *B. cinerea* resistance while knockdown of either gene alone does not (Cantu *et al*., 2008). Regardless, the PL enzyme is a substantial susceptibility factor in tomato fruit and targeting this enzyme for breeding purposes may improve fungal resistance in addition to lengthening shelf life by slowing the softening process.

Susceptibility and resistance to necrotrophic pathogens is ultimately a complex, multigenic trait in plants. The use of transcriptomic datasets to facilitate a systems-level approach of such pathosystems has increased in recent years (Alkan *et al*., 2015; Petrasch *et al*., 2019; Zhang *et al*., 2019; Kovalchuk *et al*., 2019) and has led to novel insights in both host and pathogen features that impact the outcome of such interactions. Moreover, the additional layer of an enormously developmental change such as ripening only further increases the need for these approaches. We have demonstrated how such an approach can yield critical information on both fruit infection response and broad ripening-associated changes that increase susceptibility, and additionally provide insights into single genes with a disparate impact on susceptibility. From our results, we believe that ripening-associated susceptibility is best explained by a dominant role of susceptibility factors that increase during ripening which, coupled with a modest loss of preformed defenses, outweighs the efforts of the defense response in ripe fruit (**Fig. 6**). Overall, our results have tremendous utility for guiding future study of fruit-pathogen interactions in addition to providing breeders with information on potentially useful genes for targeting in the hopes of ultimately reducing postharvest losses in tomatoes and other fruit crops.

**Fig. 6.**
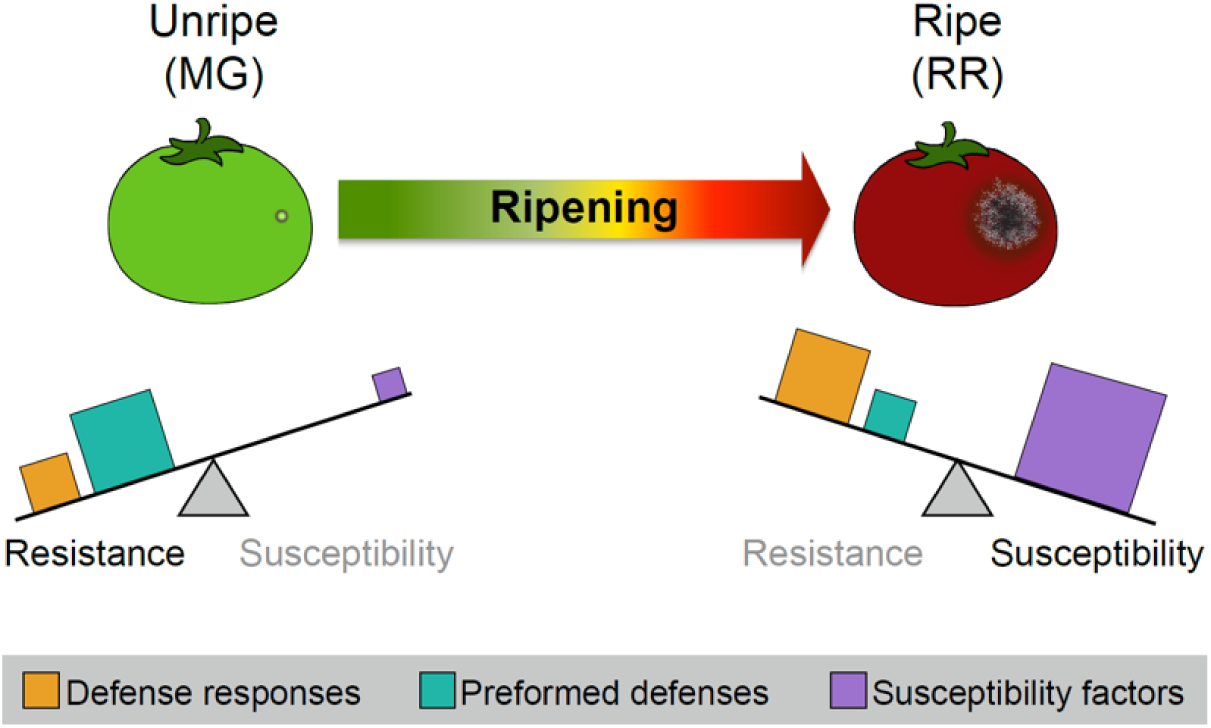
Model of contributing factors to ripening-associated susceptibility in tomato fruit. Sizes of squares indicate the relative magnitude of that feature in fruit of that stage. The balance between contributing components determines the ultimate outcome of the infection. MG = mature green, RR = red ripe.

## Supplementary Data

Fig. S1. Pathogen measurements and wound responses.

Fig. S2. Defense responses and hormone levels in wild-type and mutant fruit.

Table S1. Summaries of read mapping to tomato and pathogen transcriptomes.

Table S2. Differential expression output with functional annotations.

Table S3. Common and unique differentially expressed genes for fruit inoculated with each pathogen.

Table S4. Core RR response defense genes not expressed at equal or greater levels than MG in infected fruit.

Table S5. Enrichment of defense genes in filtered upregulated/downregulated ripening genes.

### Acknowledgments

This work was supported by start-up funds from the College of Agricultural and Environmental Sciences and the Department of Plant Sciences (UC Davis) to BB-U. Funding to CJS was partially provided by the Plant Sciences GSR Award (UC Davis). We would like to acknowledge Dr. Rosa M. Figueroa-Balderas for advice on RNAseq library preparation. We thank Saskia D. Mesquida-Pesci, Stefan Petrasch, and Dr. Ann L.T. Powell for valuable feedback on the manuscript.

